# *Gys1* antisense therapy rescues neuropathological bases of murine Lafora disease

**DOI:** 10.1101/2021.02.11.430846

**Authors:** Saija Ahonen, Silvia Nitschke, Tamar R. Grossman, Holly Kordasiewicz, Peixiang Wang, Xiaochu Zhao, Dikran R. Guisso, Sahba Kasiri, Felix Nitschke, Berge A. Minassian

**Affiliations:** Program in Genetics and Genome Biology, The Hospital for Sick Children Research Institute, Toronto, ON M5G 0A4, Canada; Division of Neurology, Department of Pediatrics, University of Texas Southwestern Medical Center, Dallas, TX 75390, USA; Department of Antisense Drug Discovery, Ionis Pharmaceuticals, Carlsbad, California, USA; Department of Biochemistry, University of Texas Southwestern Medical Center, Dallas, TX 75390, USA

**Keywords:** Lafora disease, antisense oligonucleotides, neuroinflammation, glycogen synthase, therapy

## Abstract

Lafora disease is a fatal progressive myoclonus epilepsy. At root, it is due to constant acquisition of branches that are too long in a subgroup of glycogen molecules, leading them to precipitate and accumulate into Lafora bodies, which drive a neuroinflammatory response and neurodegeneration. As a potential therapy, we aimed to downregulate glycogen synthase, the enzyme responsible for glycogen branch elongation, in the disease’s mouse models. We synthesized an antisense oligonucleotide (Gys1-ASO) that targets the mRNA of the brain-expressed *glycogen synthase 1* gene (*Gys1*). We administered Gys1-ASO by intracerebroventricular injection and analyzed the pathological hallmarks of Lafora disease, namely glycogen accumulation, Lafora body formation, and neuroinflammation. Gys1-ASO prevented Lafora body formation in young mice that had not yet formed them. In older mice that already exhibited Lafora bodies, Gys1-ASO inhibited further accumulation, markedly preventing large Lafora bodies characteristic of advanced disease. Inhibition of Lafora body formation was associated with prevention of astrogliosis and strong trends towards correction of dysregulated expression of disease immune and neuroinflammatory markers. Lafora disease manifests gradually in previously healthy teenagers. Our work provides proof of principle that an antisense oligonucleotide targeting the *GYS1* mRNA could prevent, and halt progression of, this catastrophic epilepsy.

## Introduction

Lafora disease is a teenage-onset progressive myoclonus epilepsy that results from a disturbance of glycogen metabolism. Normally, glucose chains longer than 12 units start precipitating, yet glycogen with 55,000 glucose units is soluble. The latter is achieved through actions of two enzymes, the chain-elongating glycogen synthase and glycogen branching enzyme. These enzymes act in concert to generate molecules that are highly and symmetrically branched, with branches that are short and pointing away from each other, allowing the molecules to be permeated with water and soluble.^1,2^ In recent years two additional enzymes have been shown necessary for proper glycogen structure, namely the glycogen phosphatase laforin (EPM2A) and its interacting ubiquitin E3 ligase malin (EPM2B/NHLRC1).^2^ The mechanisms by which laforin and malin regulate glycogen structure remain unclear. What is clear is that loss of either’s function results in slow but continuous generation of a portion of glycogen molecules with overlong branches, which, perhaps by winding around each other and extruding water, cause affected molecules to precipitate and over time accumulate into Lafora bodies (LBs). In the brain, by teenage years the accumulating LBs drive a progressive neuroinflammatory process, which at least in part underlies the disease’s intractable epilepsy, neurodegeneration and dementia.^2-4^

The *Epm2a*^*-/-*^ and *Epm2b*^*-/-*^ Lafora disease mouse models recapitulate the primary pathologies of the disease, namely the glycogen branch over-lengthiness, accumulation of LBs, and neuroinflammation. Downregulating glycogen synthase by breeding Lafora disease mice with mice deficient of genes for the brain-expressed glycogen synthase isoform (*Gys1*) or of its activator proteins prevents these abnormalities. This effect was achieved both by complete or partial (30-50%) reduction of brain GYS1 activity.^5-11^ Additionally, conditional knockdown of *Gys1* after disease onset, i.e. after LBs have already appeared, halts further LB formation and attenuates neuroinflammation.^12,13^

In the present work we designed an antisense oligonucleotide (ASO) that targets *Gys1* mRNA. Delivered into the CSF of Lafora disease mouse models, the ASO halted LB formation and reduced neuroinflammation. The results open a path to an ASO-based therapy for Lafora disease.

## Materials and methods

### Mice and delivery of ASOs

*Epm2a*^*-/-*^ and *Epm2b*^*-/-*^ mouse models were described previously.^5,6^ Animal procedures were approved by the Toronto Centre for Phenogenomics or Ionis Pharmaceuticals Institutional Animal Care and Use Committees. Male and female mice were anesthetized with isofluorane, and, except where indicated otherwise, 300 µg ASO in 10 µl PBS were injected intracerebroventrically (ICV) per indicated schedule, alternating ventricles at successive time points. Stereotactic injection coordinates were 0.3 mm anterior/posterior (anterior to bregma), 1.0 mm to right or left medial/lateral and -3.0 mm dorsal/ventral. Analgesic (Metacam 2 mg/kg) was administered before and for two days post injections. Gys1-ASO sequence is 5′-CATGCTTCATTTCTTTATTG-3′. Littermate controls were a no-target ASO, 5′-CCTATAGGACTATCCAGGAA-3′ (Ctrl-ASO), or PBS. Mice were sacrificed by cervical dislocation. One hemisphere was snap-frozen in liquid nitrogen for qRT-PCR and biochemical analyses, the other immersed in 10% neutral buffered formalin for histo-and immunohisto-pathology.

### Gys1 expression analysis in mouse brain samples

For most trials, RNA was extracted with the Qiagen RNeasy Lipid Tissue Mini Kit (Qiagen, #74804) and using a 1 cc syringe and 21 g needle for tissue homogenization. DNA was digested with DNase I (Thermo Scientific, #EN0521). cDNA was synthesized from 1 μg RNA using the iScript Advanced cDNA Synthesis Kit (Bio-Rad, #1725037). For the 3-12 mo trial, tissue was homogenized in Ambion TRIzol (Thermo Fisher Scientific, #15596018). Homogenate was loaded onto Phase Lock Gels (VWR, #10847-802). After centrifugation the aqueous phase was loaded onto RNA columns (PureLink™ RNA Mini Kit, Thermo Fisher Scientific, #12183025). On-column DNaseI digestion was performed using the PureLink™ DNase Set (Thermo Fisher Scientific, #12185010). cDNA was synthesized using the iScript™ Reverse Transcription Supermix kit (Bio-Rad, #1708841).

For qRT-PCR we used either the Stratagene Mx3005P (Agilent Technologies) or QuantStudio 7 Pro SmartStart (Applied Biosystems) real-time PCR system, and SYBR Green technology (PowerUp SYBR Green Master Mix [Applied Biosystems] or iTaq Universal SYBR Green Supermix [Bio-Rad]). Primer sequences were Gys1-F: 5’-CGCAAACAACTATGGGACAC-3’, Gys1-R: 5’-AGCAAGCAATGACAGGGAAG-3’, C3-F: 5’-CTGTGTGGGTGGATGTGAAG-3’, and C3-R: 5’-TCCTGAGTGTCGTTTGTTGC-3’. *Gapdh* served as reference, except for the 3-12 mo trial where *Hprt* and *Rpl4* were used. ΔCt values were determined, calculating Ct_gene of interest_ -Ct_reference gene_ (geometric mean was used in case of two reference genes), followed by transformation into 2^-ΔCt^. Expression levels were further normalized to the PBS or Baseline group.

### Protein and glycogen analyses

Frozen tissue was homogenized with buffer including Pierce protease and phosphatase inhibitors (Thermo Scientific) and 2 mM DTT. Protein concentrations were determined using the DC Protein Assay (BioRad). Equal protein amounts (30 µg) were heated in sample buffer (70°C, 10 min) and loaded to 10% SDS-PAGE for Western blotting. GYS1 and GAPDH antibodies were from Cell Signaling (#3886) and Santa Cruz (sc-365062), respectively. Immunoblots were detected with HRP-conjugated secondary antibodies and Clarity Western ECL-substrate (BioRad) and analyzed with the ChemiDoc Imaging system (BioRad). Glycogen extraction, amyloglucosidase digestion and glucose determination were as previously described.^14^

### Histological analyses

Formalin-fixed paraffin-embedded brain tissues were sectioned and stained using periodic acid-Schiff diastase (PASD) for LBs^9^ or immunohistochemistry against GFAP (mouse anti-GFAP, BioGenex, #AM020-5M; dilution 1:250). Slides were scanned using Pannoramic (3DHistech) or NanoZoomer 2.0-HT (Hamamatsu Photonics) scanners. LB and GFAP signals were quantified in the hippocampus using HistoQuant (3DHistech) by defining LBs or GFAP signals based on pixel color. Values are expressed as % area.

### Lafora body size distribution

Per hippocampal section, HistoQuant analysis yielded in a list of individual LB sizes). LBs were assigned to 30 size bins with limits defined through a 3^rd^ order binomial function: lower limit (n) = 0.00222(n-1)^3^ + 0.002; upper limit (n) = 0.00222n^3^ + 0.002; n = bin number. The number of bodies per bin was divided by hippocampal area and expressed as area-normalized LB number (n/µm^2^). The average area-normalized body number per bin was calculated across all hippocampi of each group (n>7, biological replicates) and plotted against the bin center (average of lower and upper size bin limits). To calculate fold-change from baseline, area-normalized counts of each bin were divided by the average area-normalized count of the baseline group in the same bin (also performed with individual baseline values to obtain fold-changes to baseline average for each animal). Average fold-change and SEM were plotted against body size (bin center). Statistical analyses were performed comparing group average area-normalized body numbers and fold-changes for each bin.

### Statistical analyses

Data are presented as mean ± SEM. Statistics were performed with GraphPad Prism 8.4.3. If not stated otherwise, one-way ANOVA was performed, followed by post-hoc tests with Tukey multiple comparison correction. For Fig. 3A and B and Fig. 4E two-way ANOVA was performed, also followed by post-hoc tests with Tukey multiple comparison correction. For Fig. 3D and E one-way ANOVA was performed, followed by post-hoc tests with Holm-Bonferroni multiple comparison correction. For qRT-PCR data in Fig. 2 and Fig. 4 Kruskal-Wallis ANOVA was used, followed by two-stage linear step-up procedure of Benjamini, Krieger and Yekutieli. Asterisks denote statistical significance as follows: *, p < 0.05; **, p < 0.01; ***, p < 0.001; ****, p < 0.0001.

### Data availability

The raw data that support the findings of this study are available from the corresponding authors, upon request.

## Results

### Identification of an active ASO

We designed 300 ASOs targeting different regions of *Gys1* pre-mRNA, which we screened for efficacy in mRNA downregulation. We selected Gys1-ASO for its potent (75%), yet not complete, *Gys1* mRNA downregulation (Supplementary Fig. 1A). The IC_50_ of Gys1-ASO in cell-based experiments was 3.4 µM (Supplementary Fig. 1A and B). We administered Gys1-ASO into ventricles of 1-month-old (mo) wild-type (WT) mice, sacrificed the mice 14 days later and measured *Gys1* mRNA, which had diminished by greater than 50% in the cortex, hippocampus and spinal cord (Supplementary Fig. 1C-E).

### Gys1-ASO prevents LB formation in young mice

In the Lafora disease mouse models, LBs begin to form around 1 mo of age and become increasingly visible over the following two months, especially in the hippocampus. We delivered Gys1-ASO by ICV injection to *Epm2a*^*-/-*^ mice at 1 and 2 mo and sacrificed the mice at 3 mo of age. Both mRNA and protein levels of *Gys1* were reduced by greater than 80% in the Gys1-ASO-treated group (Fig. 1A-C). Brain glycogen levels are elevated in Lafora disease mice due to the accumulation of malstructured glycogen in the form of LBs.^14^ In Gys1-ASO-treated mice brain glycogen levels were more than 50% lower than in control mice (Fig. 1D) and comparable to levels normally seen in WT mice (approximately 2 µmol/g tissue fresh weight).^14^ Finally, PASD staining revealed that LBs were likewise and equivalently reduced (Fig. 1E and F). Results of these experiments in *Epm2b*^*-/-*^ mice were similar (Supplementary Fig. 2).

**Figure 1:**
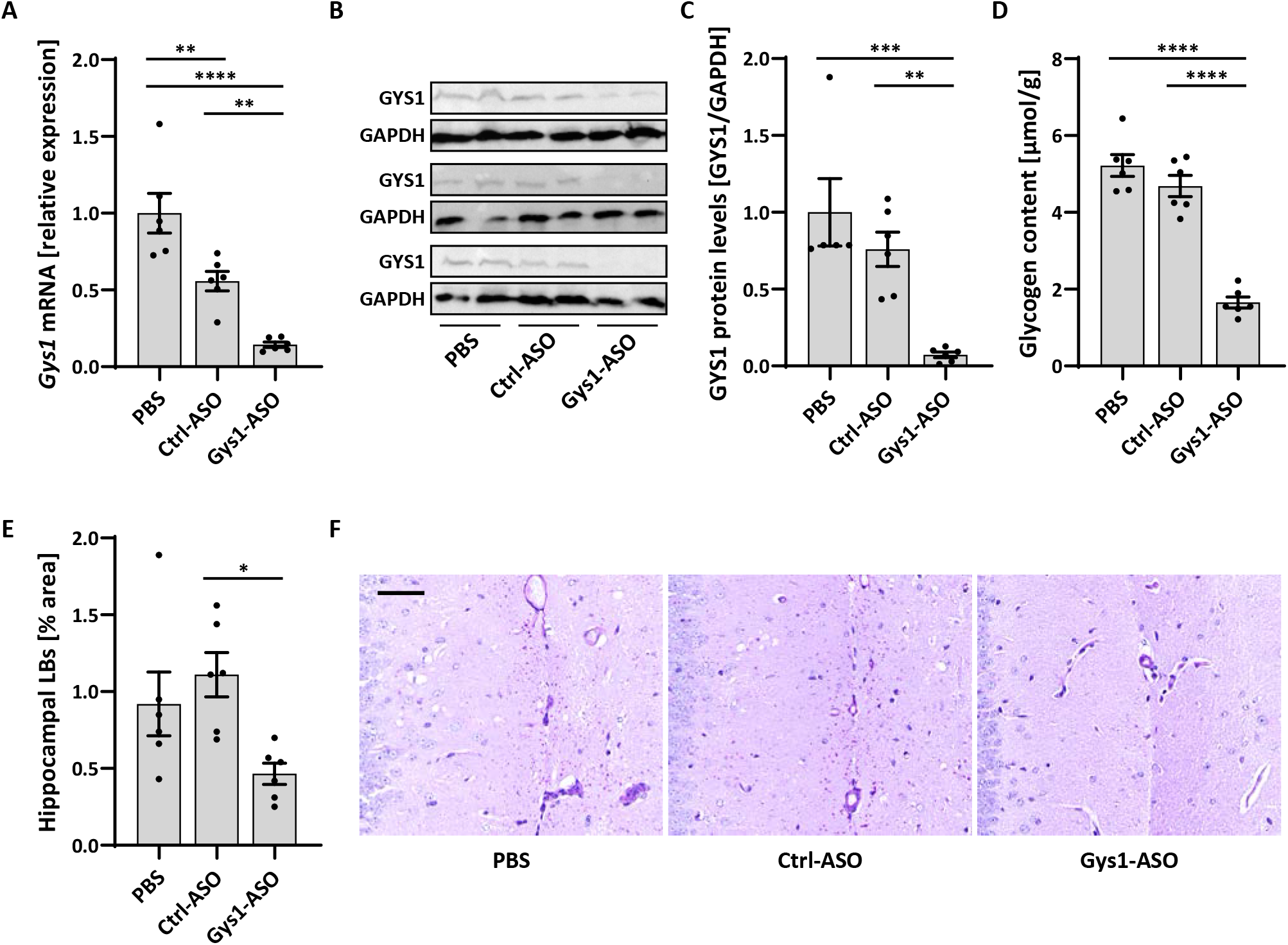
*Gys1*-targeting ASO Gys1-ASO, administered at 1 and 2 months, leads to reduced *Gys1* mRNA and GYS1 protein levels and attenuates glycogen and LB accumulation in *Epm2a*^*-/-*^ mice at 3 months. (**A**) Brain *Gys1* mRNA relative expression levels in PBS-, Ctrl-ASO-, and Gys1-ASO-injected *Epm2a*^*-/-*^ mice. Ctrl-ASO, a no-target control ASO. (**B**) Brain GYS1 Western blots with GAPDH as loading control. (**C**) Quantification of GYS1 Western blots shown in **B**, normalized to GAPDH. (**D**) Brain total glycogen content. (**E**) Lafora body (LB) quantification in the hippocampus. (**F**) Representative images of PASD stained hippocampus. Scale bar, 50 µm. All data are presented as mean ± SEM. Significance levels are indicated as *, p < 0.05; **, p < 0.01; ***, p < 0.001; ****, p < 0.0001.

The above experiments showed that when administered at the point in time when LBs are starting to appear, Gys1-ASO inhibits LB accumulation. Results being similar using both mouse models, we limited subsequent experiments to the *Epm2a*^*-/-*^ genotype.

### Gys1-ASO prevents LB formation in older mice

To determine whether the oligonucleotide is active in the context of pre-existing LB accumulations, we administered Gys1-ASO to older mice (3 mo and 8 mo old) with different specified dosing frequencies, doses and ages at sacrifice (Figs. 2 and 3A-C). These experiments were also designed in such a way as to inform us whether treatment with Gys1-ASO could reverse existing LB accumulations, i.e. not only prevent new accumulations, but lead to removal of existing ones. The results can be summarized as follows. Gys1-ASO does prevent LB accumulations beyond amounts present at time of treatment initiation. This effect is stronger when treatment starts earlier (3 mo versus 8 mo), is longer (3 mo to 12 mo versus 3 mo to 6 mo), or is dosed higher (500 µg versus 300 µg) (Figs. 2 and 3A-C). LBs do not diminish below their existing levels.

**Figure 2:**
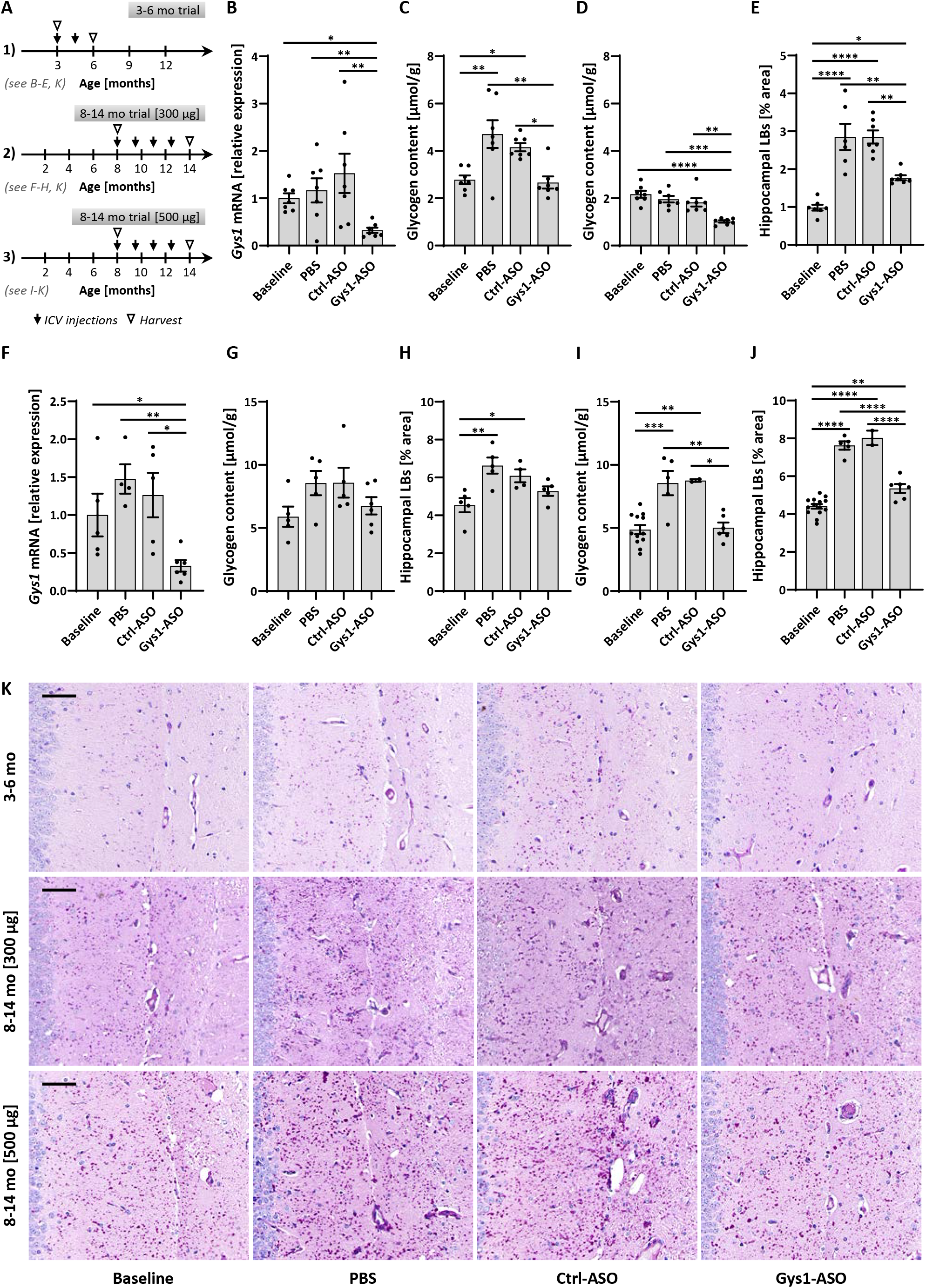
Later ASO administration, after LD onset, effectively slows disease progression in *Epm2a*^*-/-*^ mice without signs of reversal. (**A**) Experimental design of three different trials. PBS, Ctrl-ASO (no-target control ASO), or Gys1-ASO were injected and mice sacrificed at indicated time points. Untreated mice, sacrificed at time of first injection, served as baseline control. (**B** to **E**) Results from 3-6 mo trial (trial design in **A**), showing *Gys1* mRNA levels in *Epm2a*^*-/-*^ (**B**), brain total glycogen content in *Epm2a*^*-/-*^(**C**) and WT (**D**) and Lafora body (LB) quantification in the hippocampus of *Epm2a*^*-/-*^ mice (**E**). (**F** to **H**) Results from 8-14 mo trial, using 300 µg ASO for each injection (trial design in **A**), showing *Gys1* mRNA levels (**F**), brain total glycogen content (**G**) and LB quantification in the hippocampus (**H**) of *Epm2a*^*-/-*^ mice. (**I** and **J**) Results from 8-14 mo trial, using a higher dose of 500 µg ASO for each injection (trial design in **A**), showing brain total glycogen content (**I**) and LB quantification in the hippocampus (**J**) of *Epm2a*^*-/-*^ mice. (**K**) Representative images of PASD stained hippocampus of *Epm2a*^*-/-*^ mice from the three different trials, explained in **A**. Scale bar, 50 µm. All data are presented as mean ± SEM. Significance levels are indicated as *, p < 0.05; **, p < 0.01; ***, p < 0.001; ****, p < 0.0001.

**Figure 3:**
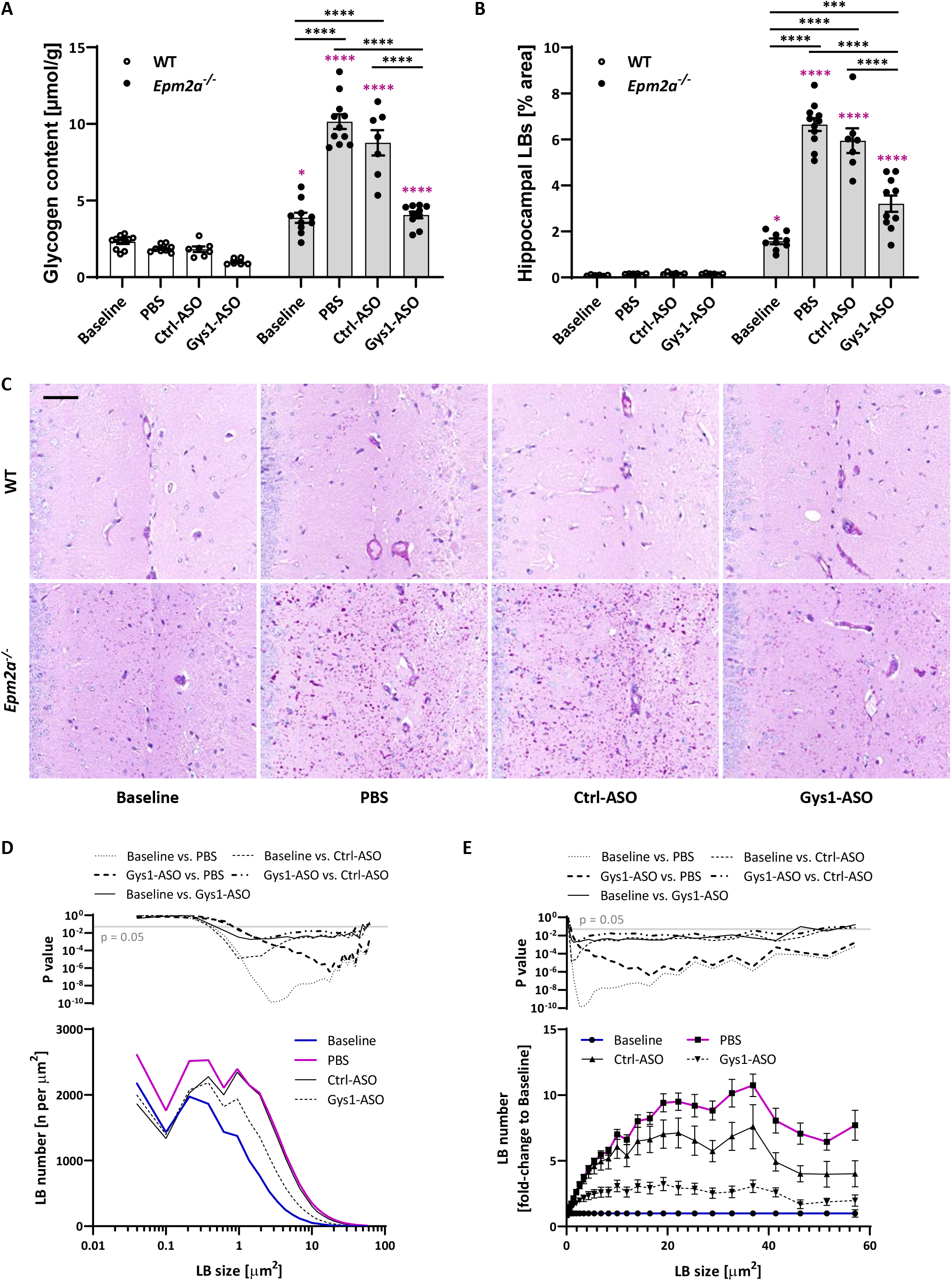
Long-term ASO treatment strongly prevents glycogen and LB accumulation in *Epm2a*^*-/-*^ mice. (**A**) Brain total glycogen content. (**B**) Lafora body (LB) quantification in the hippocampus. (**C**) Representative images of PASD stained hippocampus. Scale bar, 50 µm. PBS, Ctrl-ASO (no-target control ASO), or Gys1-ASO were injected at 3, 6, and 9 months and brain tissue analyzed at 12 months. Untreated mice, sacrificed and analyzed at 3 months, served as baseline control. Significance levels are indicated as *, p < 0.05; ***, p < 0.001; ****, p < 0.0001. Asterisks in pink indicate significance levels compared to the corresponding WT. (**D**) LB size distribution in *Epm2a*^*-/-*^ mice. (**E**) Differential LB size distribution (fold change compared to Baseline) in *Epm2a*^*-/-*^ mice. Top graph in D and E, p values comparing LB number between indicated experimental groups at different LB area bins. All data are presented as mean ± SEM.

### Gys1-ASO prevents a shift toward larger LBs

LBs occur along a spectrum between two main morphologies, small and dust-like and large and spherical. The first are much smaller and much more numerous and are located in countless astrocytic processes. The second are mostly juxtanuclear in neuronal perikarya, occupying varying extents of neuronal cytoplasms.^15,16^ We studied the size distribution of hippocampal LBs in one of our experiments (3-12 mo trial) and show that across all sizes LB abundance increases with Lafora disease progression (PBS/Ctrl-ASO versus baseline), with larger LBs (>1 µm^2^) contributing more to the overall increase of LBs (Fig. 3D and E). Gys1-ASO attenuated the growth of LBs of all sizes, in particular maintaining the abundance of larger bodies (1-60 µm^2^) low compared to controls where large bodies markedly increased in number (Fig. 3E).

### Gys1-ASO prevents Lafora disease-related astrogliosis

A remarkable 94% of genes upregulated in Lafora disease mouse model brain transcriptomes are ones involved in inflammatory and immune system pathways, strongly suggesting that immune disease at least in part underlies the disease. Of the hundreds of these genes a set of nine has been validated by qRT-PCR experiments, showing gradual increase in expression with Lafora disease progression.^4^ We measured expressions of four of these genes (*Lcn2, Cxcl10, Ccl5*, and *C3*) (as well as that of *Gys1*) in the 3-12 mo study and show that Gys1-ASO imparts a strong corrective tendency on the expression levels of all (Fig. 4A-E). We next studied whether microgliosis and astrogliosis, previously reported in murine Lafora disease, are improved. Aside from LBs, astrogliosis is the earliest and most constant neuropathological abnormality in LD mouse models.^7-10,17^ Microgliosis was not affected (Supplementary Fig. 3), but astrogliosis was corrected and near-eliminated (Fig. 4E-F).

**Figure 4:**
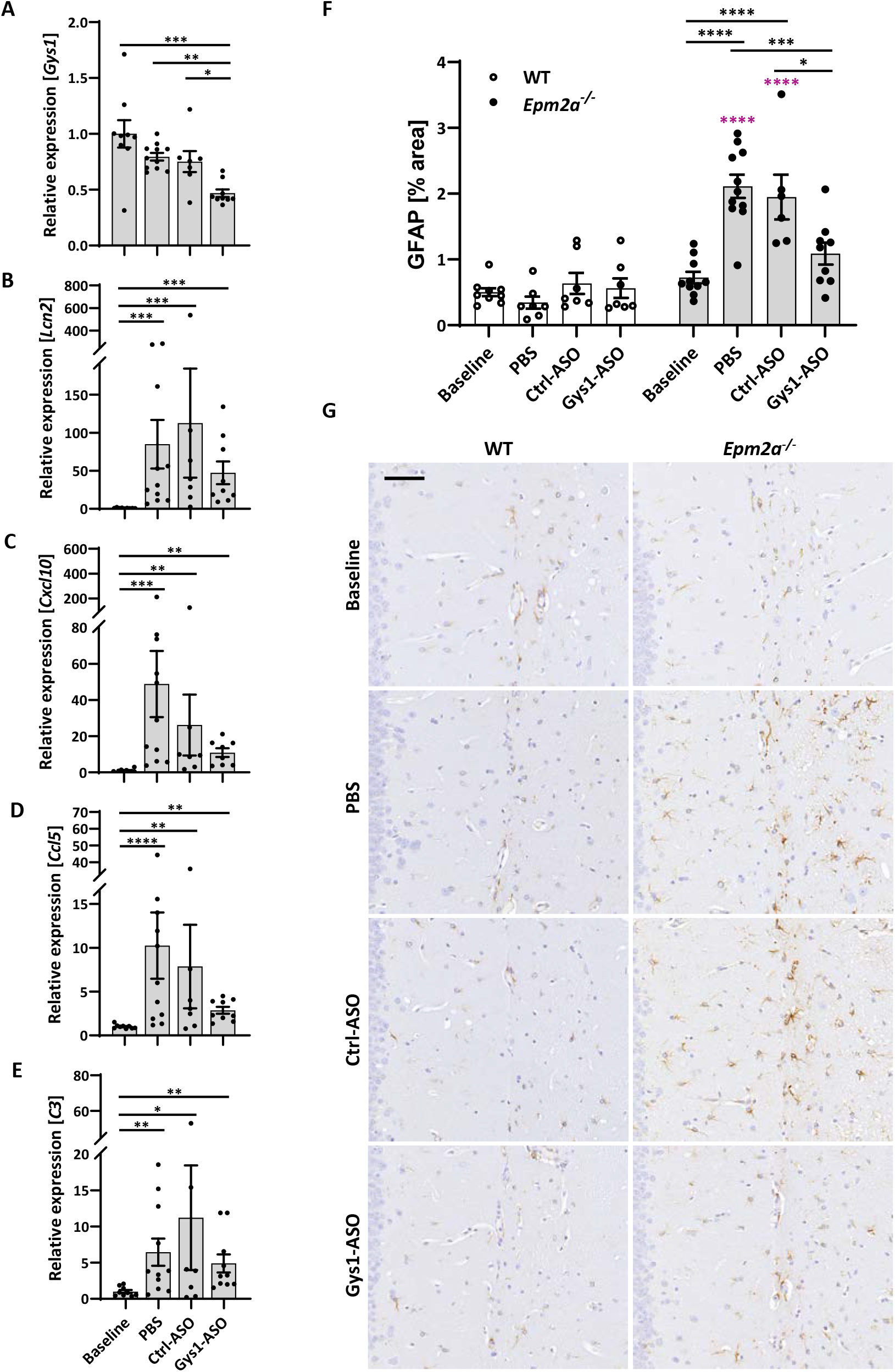
Long-term ASO treatment rescues astrogliosis in *Epm2a*^*-/-*^ mice. (**A** to **E**), mRNA relative expression levels of *Gys1* (**A**), and inflammatory and immune system response marker genes *Lcn2* (**B**), *Cxcl10* (**C**), *Ccl5* (**D**), and *C3* (**E**) analyzed by qRT-PCR. (**F**) GFAP signal quantification in the hippocampus. (**G**) Representative immunohistochemistry (IHC) images of anti-GFAP in the hippocampus. Scale bar, 50 µm. PBS, Ctrl-ASO (no-target control ASO), or Gys1-ASO were injected at 3, 6, and 9 months and brain tissue analyzed at 12 months. Untreated mice, sacrificed and analyzed at 3 months, served as baseline control. All data are presented as mean ± SEM. Significance levels are indicated as *, p < 0.05; **, p < 0.01; ***, p < 0.001; ****, p < 0.0001. Asterisks in pink indicate significance levels compared to the corresponding WT.

## Discussion

Lafora disease afflicts previously healthy children with escalating and protracted devastation. A treatment is urgently needed. While the basic mechanisms of disease are not fully elucidated, the principal cog has recently become known, namely abnormally long branches, generated by glycogen synthase, leading to glycogen insolubility.^3,18^ASOs have exceptional target specificities through Watson-Crick nucleotide sequence matching. Delivered to the CSF they distribute to practically all cells of all brain regions. They are stable and active for several months between doses.^19^ They can act through multiple mechanisms, e.g. activating a silent homologue (*SMN2*) of a mutated gene (*SMN1*) in spinal muscular atrophy,^20,21^ stabilizing and augmenting the activity of the healthy allele (*SCN1A*) in Dravet syndrome,^22^ and others. However, their original and simplest mechanism is downregulation of a target mRNA through RNaseH1,^23^ which is what is required in Lafora disease with *GYS1* as target. We here show in the mouse models of Lafora disease that an ASO targeting *Gys1* mRNA prevents the pathogenic LBs from forming or proliferating, and improves and corrects immunopathological features of the disease.

Mammals possess two glycogen synthase genes, one expressed exclusively in the liver (*Gys2*), the other (*Gys1*) in all the other organs including brain.^24^ Patients completely lacking *GYS1* (glycogen storage disease type 0b) have no neurological disease but develop cardiac arrhythmias. Parents of these patients, with 50% *GYS1* activity, are completely healthy.^25-27^ A proposed therapy for Lafora disease with a *GYS1* targeting ASO would be administered by lumbar puncture and thus not affect cardiac *GYS1*, as ASOs do not cross the blood-brain or arachnoid granulation barriers.^28-30^ In the brain, partial downregulation of *GYS1* would be aimed for and sufficient. In previous studies based on genetic crosses of LD mice with mice deficient of GYS1 or proteins that activate GYS1, we and others showed that 30-50% GYS1 downregulation suffices to prevent LB formation.^9,10^.

In the present work the ASO was effective at any disease stage, but more so in early disease and with sustained therapy. In the current era, LD patients can be diagnosed within weeks of symptoms. Myoclonic, visual or convulsive seizures lead to an EEG, which beyond epileptiform discharges shows background dysregulation and raises suspicion of a non-benign epilepsy. Since *EPM2A* and *EPM2B* are now widely included in epilepsy gene panels, diagnosis should then be swiftly made. In early stages of disease, LD patients are no different than other teenagers with new-onset epilepsy. As such, a therapy that halts the disease would be, in principle, tantamount to a cure.

Precisely how the laforin glycogen phosphatase and malin ubiquitin E3 ligase cooperate to fine-tune the lengths of glycogen branches to generate perfectly spherical and soluble macromolecules awaits to be uncovered. Meanwhile it is hoped that the present results will rapidly translate to a therapy for LD.

## Acknowledgments

We would like to thank Jennifer P. Lee and Julia Gliwa for expert technical support.

## Supplementary material

Supplementary material is available at Brain online.

## Abbreviations

GYS1: (glycogen synthase 1),
ASO: (antisense oligonucleotide),
Ctrl-ASO: (control ASO),
ICV: (intracerebroventricular),
LB: (Lafora body),
PBS: (phosphate-buffered saline),
PASD: (periodic acid- Schiff diastase),
SEM: (standard error of mean),
IC50: (half maximal inhibitory concentration),
mo: (month)

