## Supplementary Materials for "*Gys1* antisense therapy rescues neuropathological bases of murine Lafora disease"

**Running head:**

***Gys1* ASO therapy for Lafora disease**

**Saija Ahonen[1†], Silvia Nitschke[1,2,†], Tamar R. Grossman[3], Holly Kordasiewicz[3], Peixiang Wang[1], Xiaochu Zhao[1], Dikran R. Guisso[2], Sahba Kasiri[2], Felix Nitschke[2,4], Berge A. Minassian[1,2*]**

### SUPPLEMENTARY MATERIALS

**DETAILED METHODS Supplementary Figure 1:** 2′-O-methoxyethyl modified antisense phosphorothioate oligonucleotides (2′-MOE ASOs) were synthesized and purified as described previously[1,2] and dissolved in PBS (Ca-Mg-; Invitrogen) as 5-10-5 gapmers. For screening 700 µM ASOs were electroporated into B16-F10 cells using the HT-200 BTX Electroporator system in 96-well plates (BTX, 2 mm; Harvard Apparatus). Cells were harvested 16 hours post-treatment and lysed, and total RNA was extracted using RNeasy columns (Qiagen). Five different concentrations of the top ASO (henceforth Gys1-ASO) were tested in the same fashion to determine its dose response relationship and $IC_{50}$ value. *Gys1* mRNA was quantified using quantitative RT-PCR (qRT-PCR) (TaqMan) and an ABI Prism 7700 sequence detector (Applied Biosystems), with primers and probe as follows, forward: 5’-TGATGAAGAGAGCCATCTTTGC-3’, reverse: 5’-AGGAGTCGTCCAGCATGTTGT-3’, and probe: 5’-Fam-ACTCAGCGGCAGTCTTTCCCACCA-Tamra-3’. PCR results were normalized to total RNA measured by Quant-iT RiboGreen RNA Reagent (Molecular Probes) and expressed as percent of results from untransfected controls (% UTC). The same *Gys1* primer/probe set was used for central nervous system region extracts normalizing against *Cyclophilin A* (forward: 5’-TCGCCGCTTGCTGCA-3’, reverse: 5’-ATCGGCCGTGATGTCGA-3’, probe: 5’-Fam-CCATGGTCAACCCCACCGTGTTC-Tamra-3’).

**DETAILED METHODS Supplementary Figure 3:** Formalin-fixed paraffin-embedded brain tissues were sectioned and stained using immunohistochemistry against IBA1 (rabbit anti-IBA1, Wako, #01919741; dilution 1:1400). Slides were scanned using the NanoZoomer 2.0-HT (Hamamatsu Photonics) slide scanner. IBA1 signals were quantified in the hippocampus using HistoQuant (3DHistech) by defining IBA1 signals based on pixel color. Values are expressed as % area.

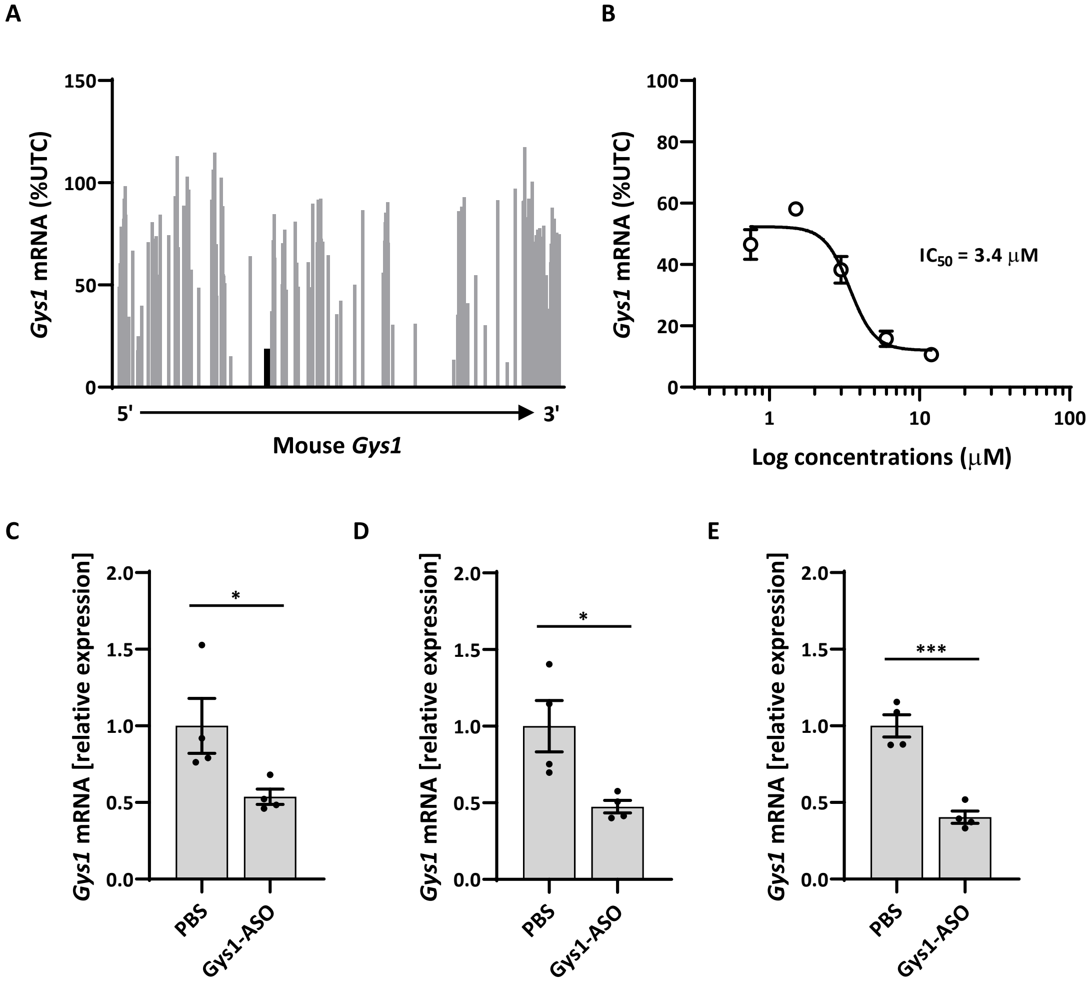


**Supplementary Fig. 1: Selection of a *Gys1*-targeting ASO for further mouse studies.** (**A**) Screen of ASOs complementary to mouse *Gys1* in B16-F10 cells. *Gys1* mRNA levels, expressed as percent of untransfected control (% UTC). ASOs listed in order of relative binding site on mouse *Gys1* transcript (5' to 3'). One ASO (henceforth *Gys1*-ASO) was selected for further characterization (black bar). (**B**) Dose response of *Gys1*-ASO in B16-F10 cells. IC50 calculated from five-point non-linear fit dose-response curve. (**C** to **E**) *Gys1* mRNA relative expression levels in the cortex (**C**), hippocampus (**D**), and spinal cord (**E**) of PBS- or *Gys1*-ASO-injected mice. All data are presented as mean ± SEM. Significance levels are indicated as *, $p < 0.05$; ***, $p < 0.001$.

**Supplementary Fig. 1**

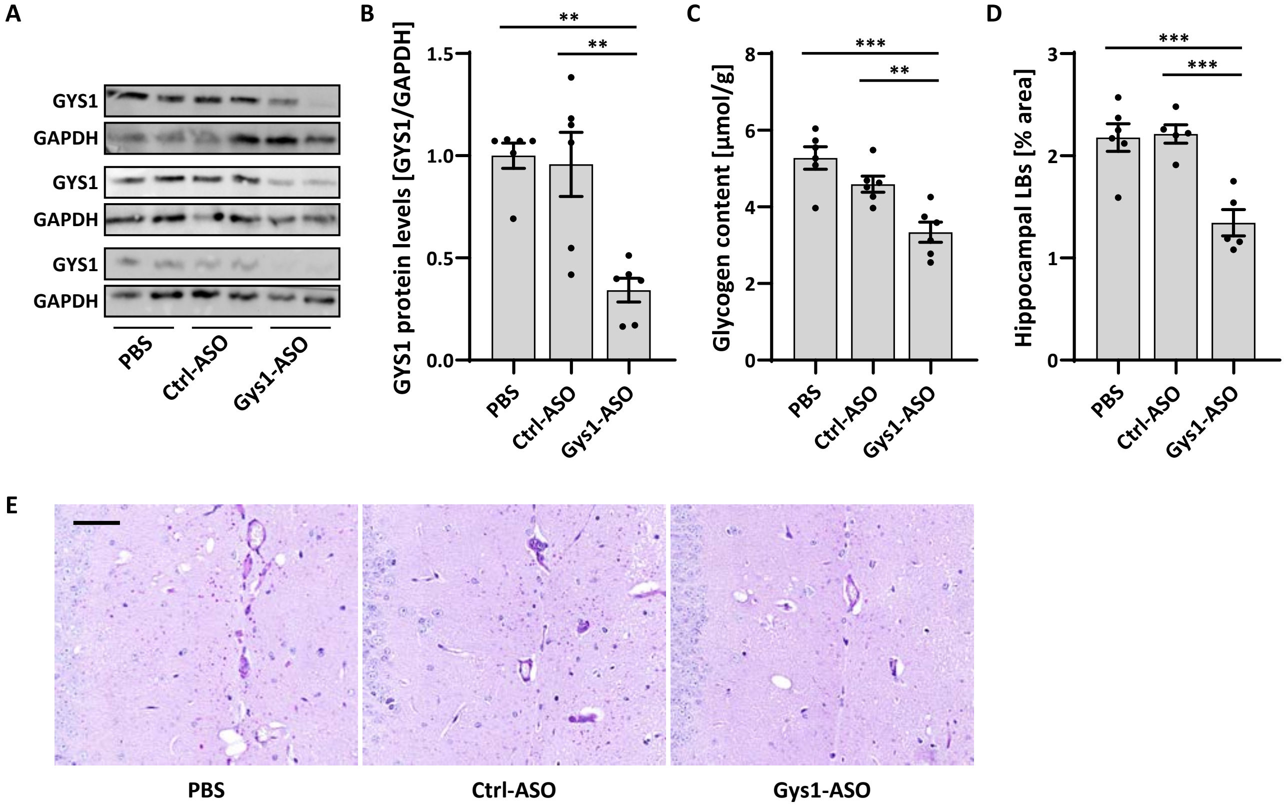


**Supplementary Fig. 2: *Gys1*-targeting ASO Gys1-ASO, administered at 1 and 2 months, leads to reduced GYS1 protein levels and attenuates glycogen and LB accumulation in *Epm2b*$^{-/-}$ mice at 3 months.** (**A**) Brain GYS1 Western blots in PBS-, Ctrl-ASO-, and Gys1-ASO-injected *Epm2b*$^{-/-}$ mice with GAPDH as loading control. Ctrl-ASO, a no-target control ASO. (**B**) Quantification of GYS1 Western blots shown in **A**, normalized to GAPDH. (**C**) Brain total glycogen content. (**D**) Lafora body (LB) quantification in the hippocampus. (**E**) Representative images of PASD stained hippocampus. Scale bar, 50 µm. All data are presented as mean ± SEM. Significance levels are indicated as **, $p < 0.01$; ***, $p < 0.001$.

### Supplementary Fig. 2

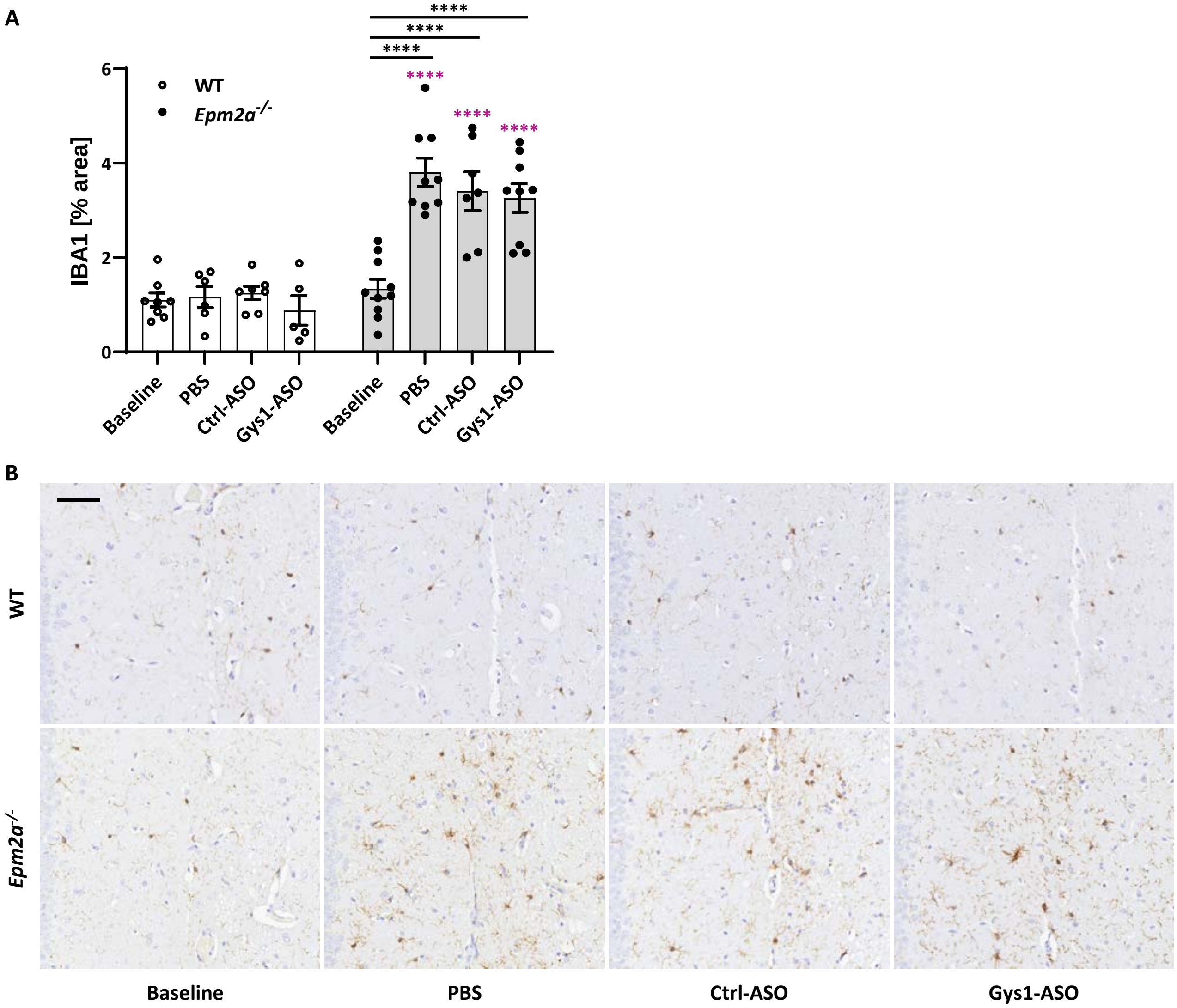


**Supplementary Fig. 3: Long-term ASO treatment does not rescue microglia activation.** (**A**) IBA1 signal quantification in the hippocampus. (**B**) Representative immunohistochemistry (IHC) images of anti-IBA1 IHC in the hippocampus. Scale bar, 50 µm. PBS, Ctrl-ASO (no-target control ASO), or Gys1-ASO were injected at 3, 6, and 9 months and brain tissue analyzed at 12 months. Untreated mice, sacrificed and analyzed at 3 months, served as baseline control. All data are presented as mean ± SEM. Significance levels are indicated as ****, $p < 0.0001$. Asterisks in pink indicate significance levels compared to the corresponding WT.

### Supplementary Fig. 3
